## Supplementary figures for "Controlling spatiotemporal pattern formation in a concentration gradient with a synthetic toggle switch"

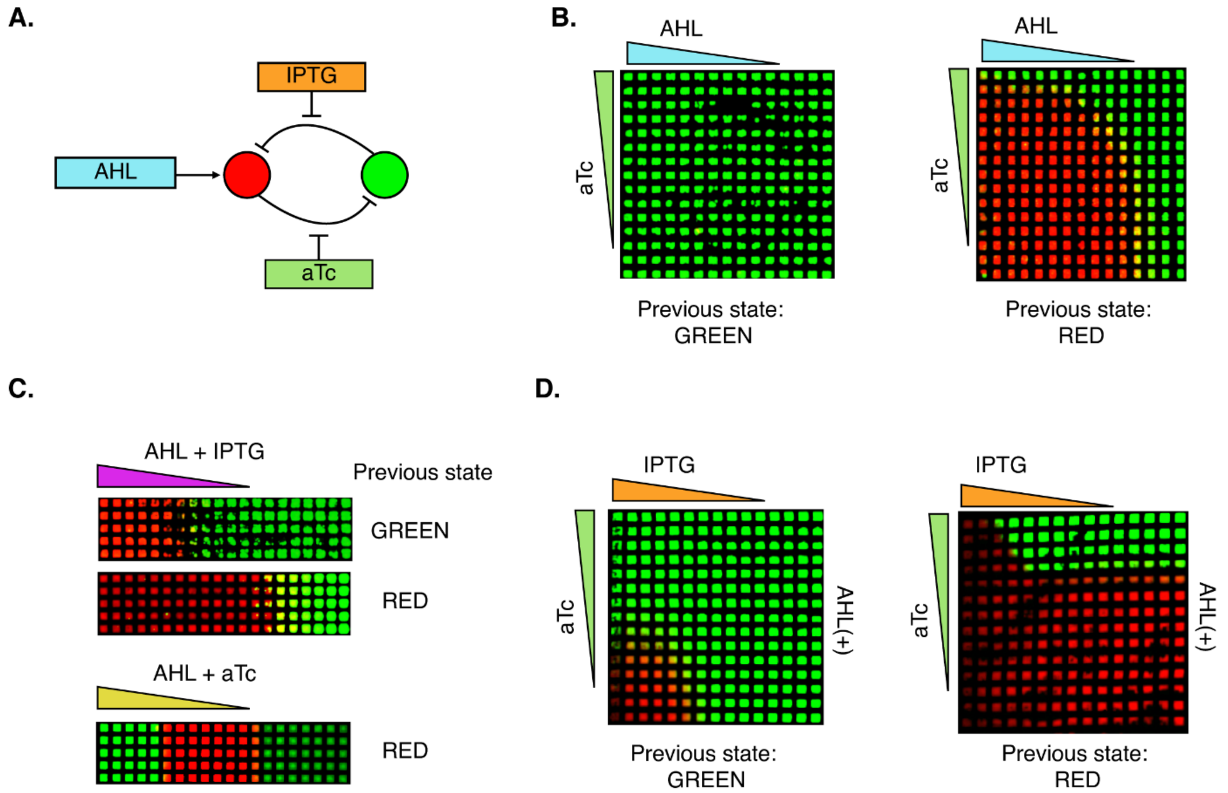


**Supplementary Figure 1: Exploration of different patterns using the inducible toggle switch**

**A**. Topology of the toggle switch network as described in Figure 1A with the addition of aTc that regulates the repression strength of the red node. **B.** Grid pattern of the circuit with aTc (0.5 ng/ml) and AHL (100 μM). If the cells were previously in the green state, the full grid stayed green, because, as shown before, IPTG is required to switch from the green to the red state. Starting from the red state, the state was conserved in the presence of AHL, but at high aTc concentrations the cells switched to the green state even in presence of high AHL concentrations. **C.** Patterning with parallel gradients. The inducer and the regulator were added together at the same edge of the grid, thus diffusing in parallel. This allows to explore the grid pattern in Figure 1 and in B in the diagonal. **D.** Patterning observed by combinations of IPTG, aTc and AHL. 100mM IPTG and 5ng/ml aTc were added as indicated and 10μM of AHL was homogenously added in the agar plate.

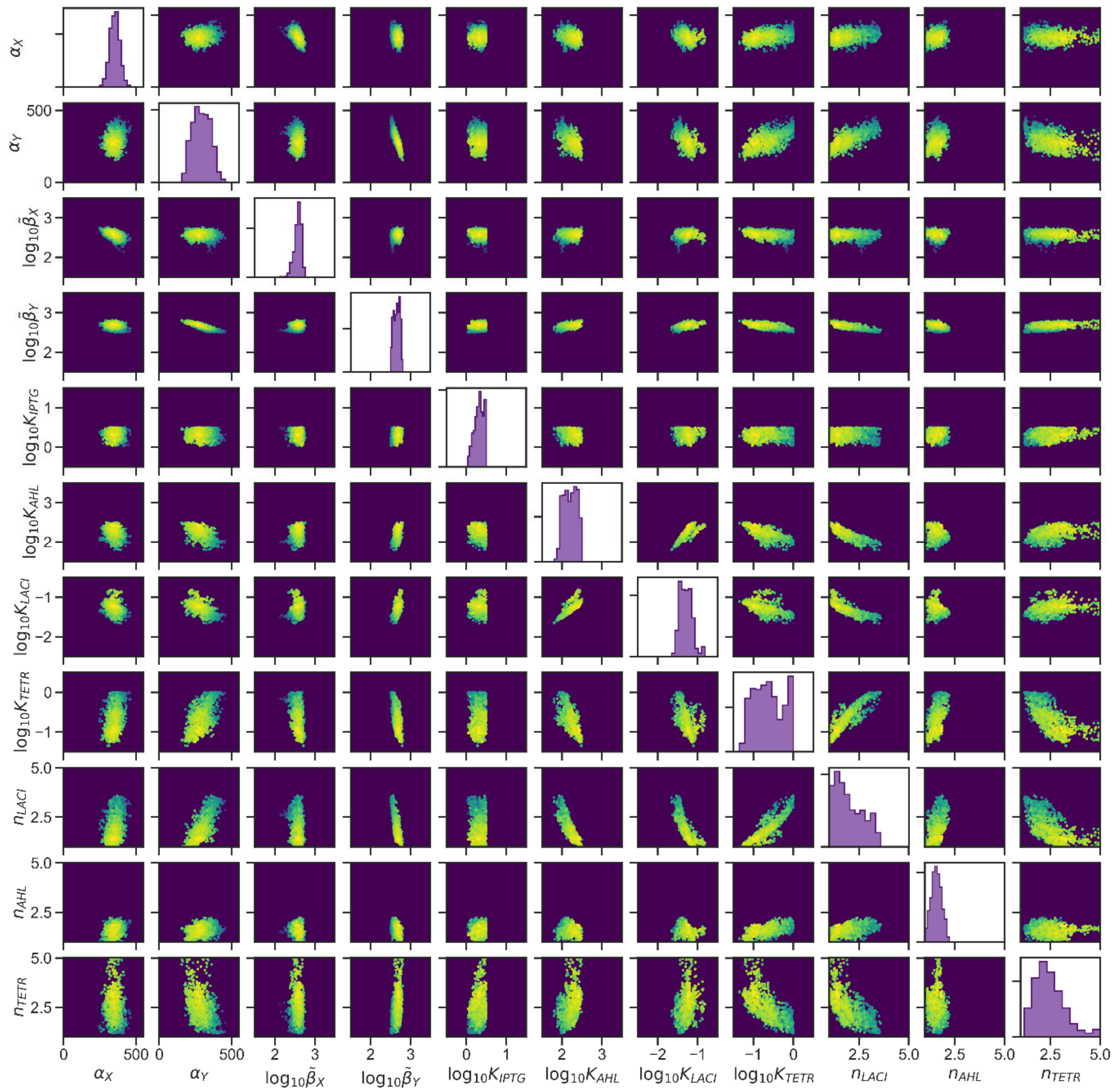


**Supplementary Figure 2: Probability space of parameters**

Result from the MCMC fitting to the flow cytometry data. Diagonal show posterior marginal probability distributions, off-diagonal show the posterior pair-wise probability distributions.

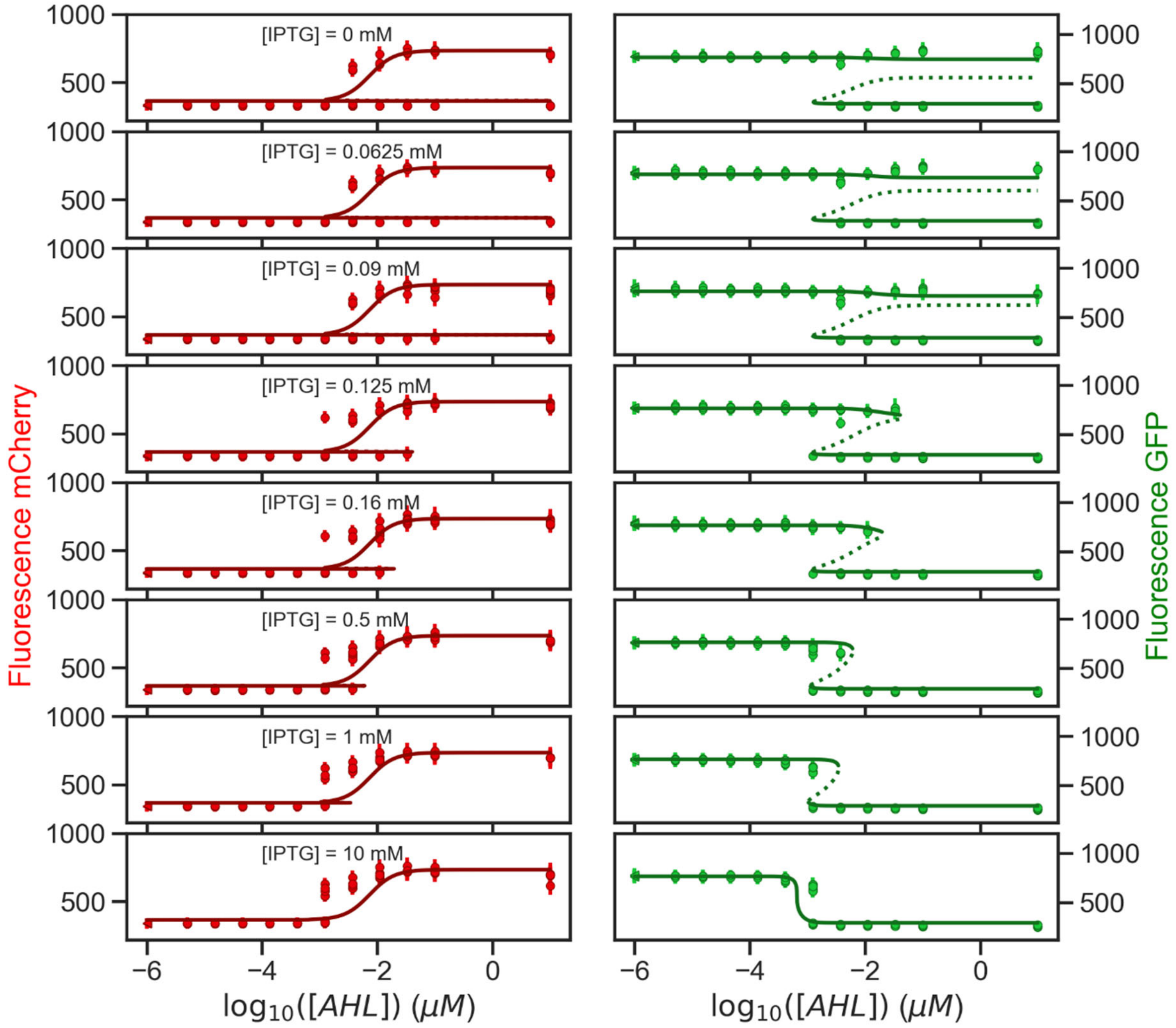


**Supplementary Figure 3: Parametrization of the mathematical model.**

Whole data set belonging to Figure 2B. Comparison between the observed populated states from the flow cytometry data (circles) and the available steady states predicted by the model (solid lines: stable states, dotted lines: unstable states). Experimentally observed states show the median and standard deviation for the different gated populations. Parameters used in the model are the best parameter candidates from the MCMC fitting.

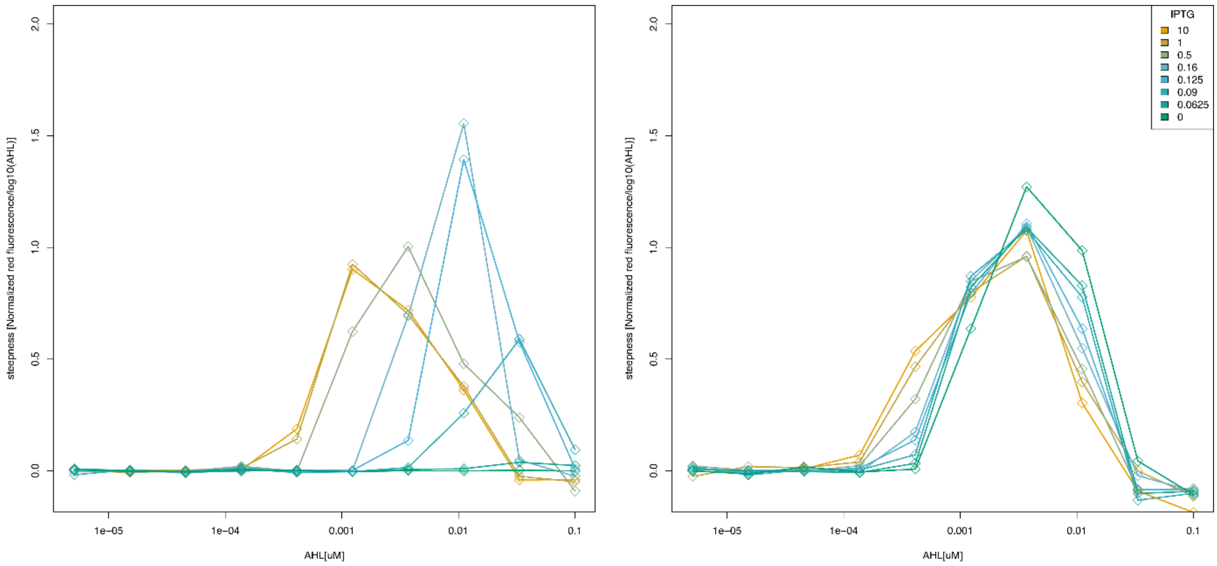


**Supplementary Figure 4: Effect of IPTG on the sharpness of the boundary**

The sharpness was quantified as the difference in red fluorescence between two consecutive points the measurements shown in Figure 2D. The higher and narrower a peak is, the sharper is the transition into the other state.

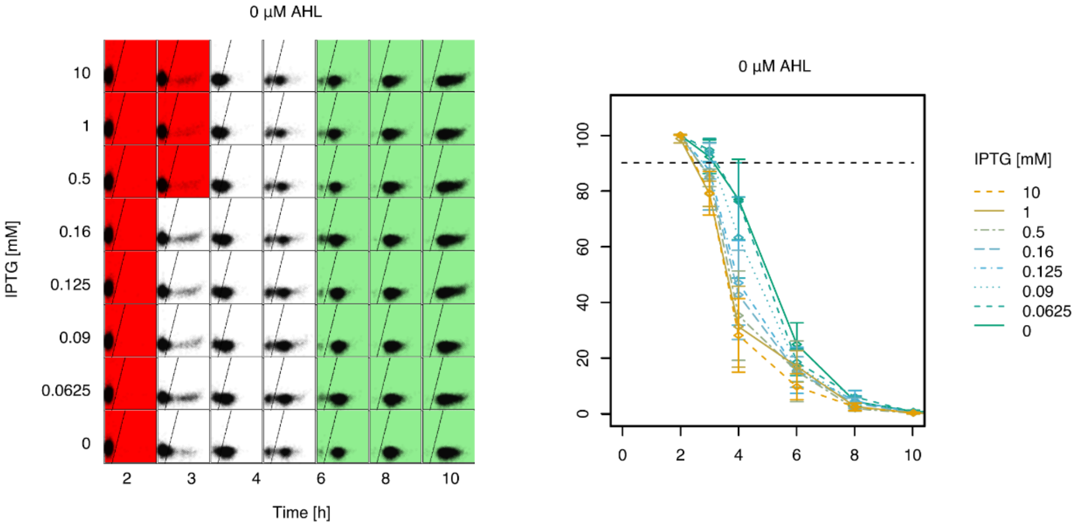


**Supplementary Figure 5: IPTG effect on the switching time of the red to the green state**

**A,B.** Left: Effects of IPTG concentration on the switching time from the red to the green state. Each square represents flow cytometer data of 10,000 events red (Y-axis) and green fluorescence (X-axis). The background color of the square indicate whether >90% of the events are in the red or the green gate. Right: Percentages of cells in the red gate over time. Mean and standard deviation of 3 biological replicates. The dotted lines represent the 90% threshold for the red color.

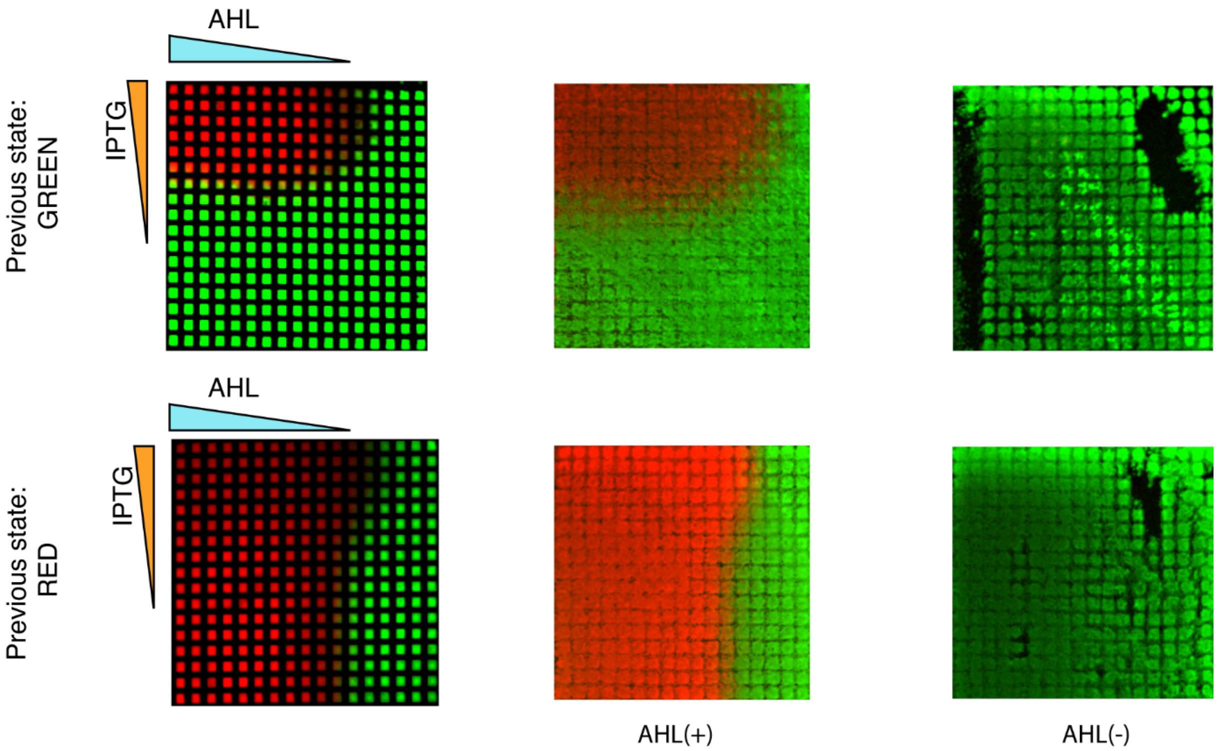


**Supplementary Figure 6: Memory propriety of the toggle switch**

Demonstration of the memory property of the inducible toggle switch. Right: grid patterning reproduced as described in Figure 1. Middle: the cells of the left grid were transferred with a paper stamp onto a new grid placed on top of an agar plate supplemented with 5 μM of AHL. We observed that the pattern was maintained. Right**:** the cells of the left grid were transferred with a paper stamp onto a new grid placed on top of an agar plate without any IPTG or AHL. The pattern was lost (control). All images were recorded after overnight incubation at 37 °C, triangles indicate the diffusion direction of the added molecules and colors represent the presence of GFP (green) and mCherry (red).

**Supplementary Movie 1: Time lapse of the flow cytometry data**

Each square represents flow cytometer data of 10,000 events red (Y-axis) and green fluorescence (X-axis). The background color of the square indicates whether >90% of the events are in the red or the green gate. From top to bottom are different concentrations of IPTG and from left to right are different concentrations of AHL as indicated in Figure 2A.
